# SodaPop: A computational suite for simulating the dynamics of asexual populations

**DOI:** 10.1101/189142

**Authors:** Louis Gauthier, Rémicia Di Franco, Adrian W.R. Serohijos

## Abstract

**Motivation:** Simulating protein evolution with realistic constraints from population genetics is essential in addressing problems in molecular evolution, from understanding the forces shaping the evolutionary landscape to the clinical challenges of antibiotic resistance, viral evolution and cancer.

**Results:** To address this need, we present SodaPop, a new forward-time simulator of large asexual populations aimed at studying their structure, dynamics and the distribution of fitness effects with flexible assumptions on the fitness landscape. SodaPop integrates biochemical and biophysical properties in a cell-based, object-oriented framework and provides an efficient, open-source toolkit for performing large-scale simulations of protein evolution.

**Availability and implementation:** Source code and binaries are freely available at https://github.com/louisgt/SodaPop under the GNU GPLv3 license. The software is implemented in C++ and supported on Linux, Mac OS/X and Windows.

**Supplementary information:** Supplementary information is available on the Github project page.

## Introduction

Evolution predominantly depends on two causalities - population dynamics and the distribution of fitness effects (DFE) (Eyre-Walker and Keightley, 2007). Despite the efforts to combine these two causalities (DePristo *et al*., 2005; Silander *et al*., 2007; Goldstein, 2011; Liberles *et al*., 2012; Goldstein 2013; Serohijos and Shakhnovich, 2016; Echave and Wilke, 2017) there remains a broad divide between population genetics and protein biophysics, both conceptually and methodically. Bridging this gap is an essential step towards our understanding of molecular evolution as a multi-scale process.

An important tool to study molecular evolution and compare outcomes of different evolutionary scenarios is simulation. Methods to perform forward simulations vary in scope and flexibility and are generally designed to investigate variety of problems in evolutionary biology and population genetics such as polymorphism and population structure (Peng and Kimmel, 2005; Padhukasahasram *et al*., 2008; Hernandez, 2008; Carvajal-Rodriguez, 2008; O’Fallon, 2010; Thornton, 2014). These programs commonly implement features such as linkage and recombination, specific migration, growth or mating schemes and selection regimes. Notably, softwares such as OncoSimulR (Diaz-Uriarte, 2017) model the evolution of large asexual populations, yet enforce strictly biallelic loci on limited sites. Likewise, there are several tools intended to model protein evolution (Pang *et al*., 2005; Blackburne and Hirst, 2005; Koestler *et al*., 2012; Grahnen and Liberles, 2012; Arenas *et al*., 2013). However, these programs are typically aimed at phylogenetic reconstruction and alignment methods testing (Ziheng and Rannala, 2012).

Regardless of the practicality of current simulation packages in addressing problems in human and population genetics, very few programs explicitly account for the DFE of proteins or integrate multiple scales in their evolutionary framework. Mutation and selection are indeed separated by increasingly complex levels of biological organization. Studying molecular evolution also requires accounting for higher-order scales such as systems and population. Moreover, most molecular evolution simulators enforce a monoclonal regime, which does not require the continuous tracking of an explicit population, but rather a single lineage. Despite the higher computational tractability of this approach, evolution in large populations such as bacterial colonies and malignant tumors is polyclonal, where the dynamics of segregating alleles is of critical importance (Greaves and Maley, 2012; Lenski, 2017).

Here we introduce SodaPop, an efficient forward-time, object-oriented (OOP) simulator aimed at studying the evolutionary dynamics of large-scale asexual populations with explicit genomic sequences. In this framework, the population structure and the DFE of fixed mutations can be explored simultaneously. Rather than being treated as a distribution (Haller and Messer, 2017; Kim *et al*., 2017), the DFE of arising mutations can be inputed from protein engineering methods (Kumar *et al*., 2006; Yin *et al*., 2007; Laimer *et al*., 2015; Jia *et al*., 2015) or from exhaustive mutagenesis experiments such as deep mutational scanning (Firnberg *et al*., 2014; Fowler and Fields, 2014; Bloom, 2014). Also, SodaPop allows full flexibility in defining fitness functions from biochemical/biophysical models that describe evolution of proteins. Additionally, the OOP framework provides a scaffold where further developments can be easily integrated.

To our knowledge, SodaPop is the first publicly available and open source tool to this end. The main program is implemented in C++ as a command-line tool. We also provide complementary tools to analyze and visualize simulation results. Source code, binaries and documentation can be downloaded freely from https://github.com/louisgt/SodaPop under the GNU GPLv3 license. Moreover, this software is portable on any POSIX-compliant operating system, including Linux and Mac OS/X, or on Windows using the Cygwin environment.

## Methods

SodaPop uses an adapted Wright-Fisher model with selection (Fisher, 1922; Wright, 1931). Populations are characterized by a top-down organization: cells are dynamic objects comprising a vector of genes, which are in turn defined by independent properties such as concentration or abundance, functional essentiality and thermodynamic stability. Genetic sequences evolve explicitly from one generation to the next, and can be traced back to the ancestral sequence through an identifier. This hierarchical, object-oriented cell model marks a first step towards a systems biology framework for the study of evolutionary dynamics. Generations are discrete time steps in which each cell object gives birth to a number of children drawn from a binomial distribution with mean equal to the fitness of the parent cell relative to the fitness summed over all cells (Figure 1). These children form the basis for the next generation of cells. Following the reproductive phase, the new population is scaled up or down to match the initial population size. This process is akin to a serial passaging bottleneck experiments (Ebert, 1998; Gullberg *et al*., 2011).

**Figure 1.**
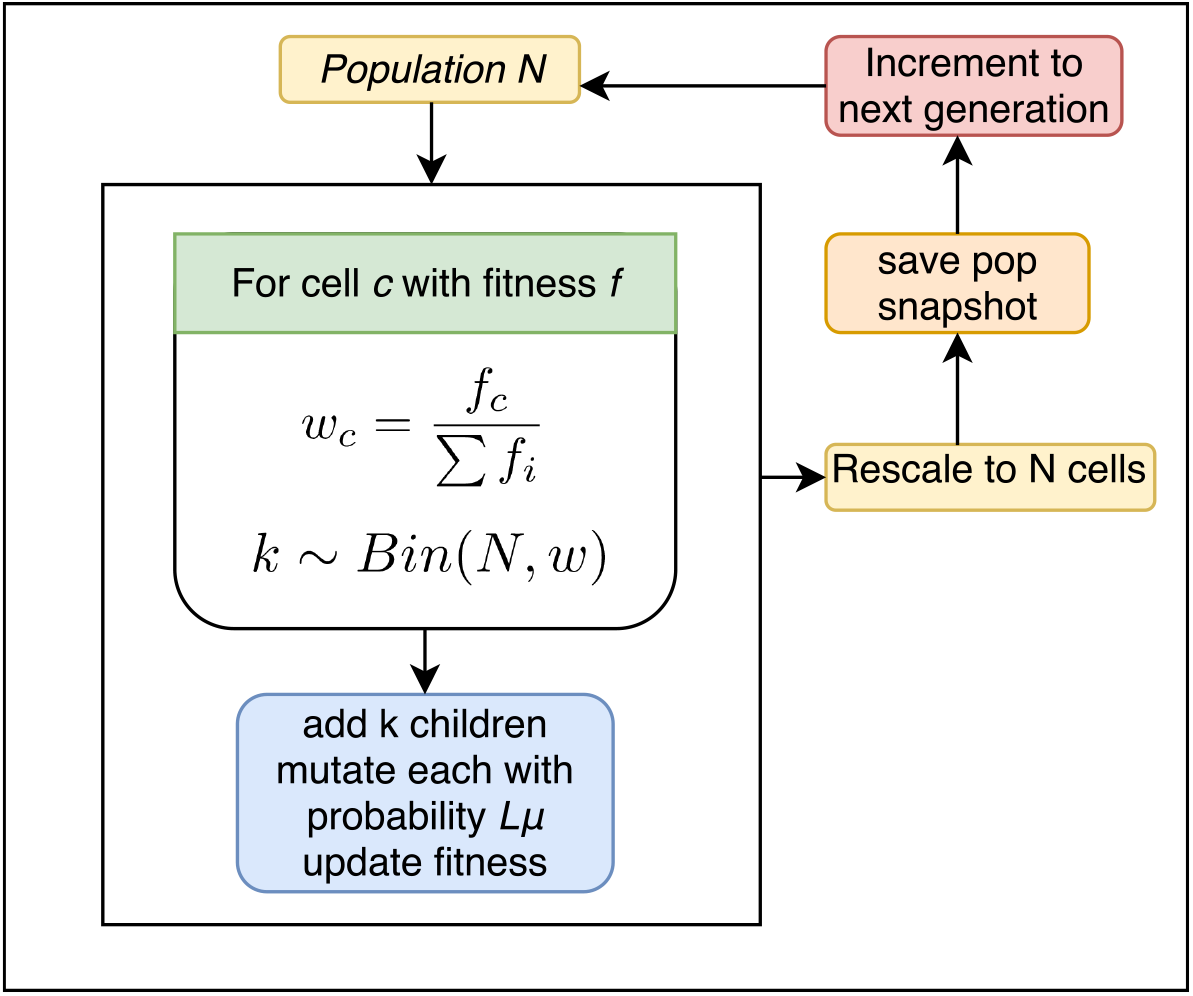
Illustration of SodaPop’s core algorithm. The Wright-Fisher process iterates through every cell and draws the number of children to add to the next generation. These children are mutated with probability *Lµ*, where *L* is the genomic length and *µ* is the mutation rate. Once the whole parent population has been swapped with daughter cells, this new generation is rescaled to *N* cells.

The program can track all arising mutations during a simulation run to provide the full history of genetic changes in the population. The program also tracks the associated selection coefficients, which enables the temporal analysis of the DFE of substitutions. In addition, SodaPop saves comprehensive snapshots of the population at a user-specified interval. This can be tuned to an arbitrary granularity to yield an explicit genealogy of sequences and analyze the clonal dynamics. The population snapshots can also be used as input for subsequent simulations. This feature facilitates the recovery of the latest state in a simulation in case of an unexpected system crash. SodaPop is built upon streamlined data structures and a fast algorithm to achieve high computational efficiency and to minimize the general trade-off between flexibility and runtime (Carvajal-Rodriguez, 2008). The program is designed to support large population sizes and rich substructures to reflect the natural magnitude of bacterial colonies and their intrinsic dynamics. As such, SodaPop can readily handle simulations in the order of 10^6^ unique cells with runtimes clocking under a few hours. We can reasonably tune the strength of selection or mutation rate to achieve higher dynamical scales without incurring a significant computational penalty.

SodaPop allows users to provide the nature and distribution of fitness effects as well as the genotype-to-phenotype relationship to use in their simulation. The DFE of arising mutations can be probabilistic, that is, defined by a distribution chosen by the user (Figure 2A). It can also be inputted from experiment (Figure 2B) or from computational estimates of biophysical properties (Figure 2C). The ability to apply a specific fitness function based on input type (Figure 2D) provides an additional layer of parameterization to the simulation. Altogether, these capabilities establish a robust framework for the investigation of theoretical and applied problems alike.

**Figure 2.**
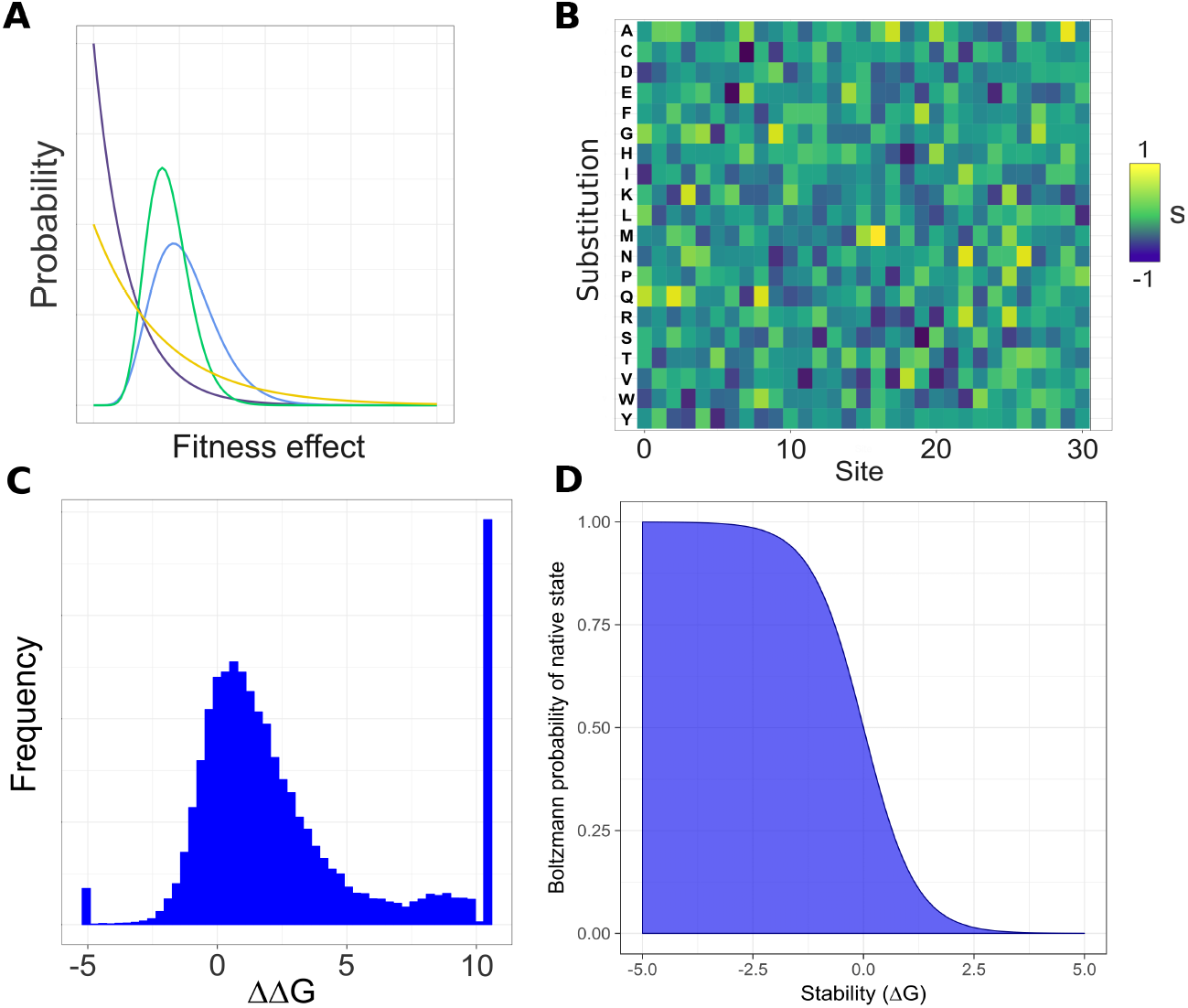
SodaPop accepts various inputs and fitness functions. **(A)** Fitness effects can be drawn from a gamma or normal distribution specified by the user. **(B)** Fitness effects may take the form of deep mutational scanning (DMS) substitution matrices for each protein, or **(C)** biophysical substitution matrices derived from computational tools. **(D)** The genotype-to-phenotype mapping is chosen by the user based on input.

## Results

In this section, we provide some examples of simulations performed by SodaPop. We present simulations of protein evolution with different population genetics parameters as well as different fitness functions.

### Test case I: Population dynamics and fitness trajectories

An evolutionary simulation with 10 genes in the folate biosynthesis pathway of *Escherichia coli* is illustrated in Figure 3. Users may also implement their own fitness function and incorporate additional protein properties such as catalytic efficiency or relative solvent accessibility (see Supporting Information for details). One of the major aims of SodaPop is to model rampant phenomena such as clonal interference and selective sweeps, which contribute significantly to population dynamics (Elena and Lenski, 2003). The ability to investigate polyclonal structure and relative fitness is of particular interest for co-culture competition assays in microbiology (Lenski *et al*., 1998; Conrad *et al*., 2011; Melnyk and Kassen, 2011; Dragosits and Mattanovich, 2013).

**Figure 3.**
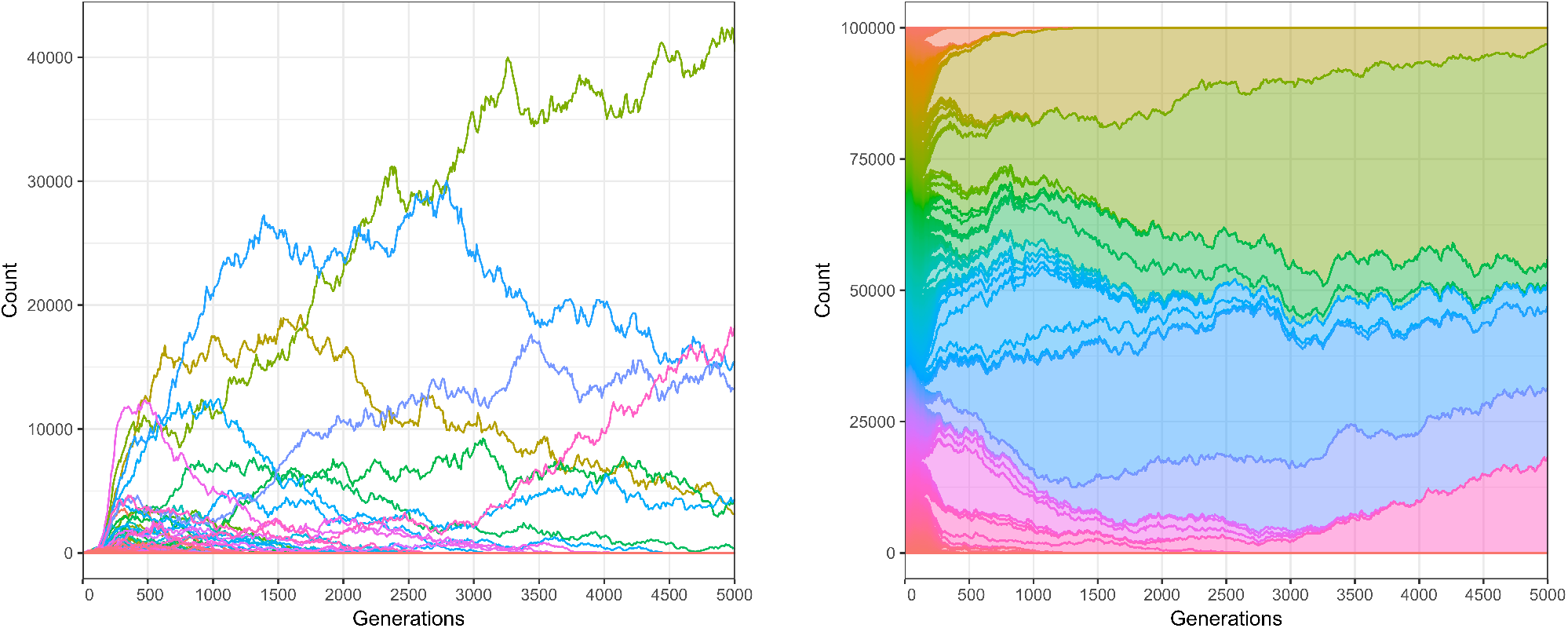
Test case 1: evolution under selection against misfolding toxicity. **(A)** SodaPop tracks the evolution of clones concurrently. Each color represents a single lineage identified by a barcode. The information in the left panel can also be represented as (B) the density of each lineage through time relative to the total population. Both these representations show pervasive clonal interference and competition.

### Test case II: Multiple sequence alignment and conservation score

To assess the performance of SodaPop in recapitulating the extent of amino acid conservation for a protein that is under selection for stability, we compared simulated protein sequences to real sequence data. It is known from *in silico* simulations that selection for protein folding stability using physical force field estimations can reproduce the pattern of sequence conservation in real biological sequences (Dokholyan and Shakhnovich, 2001; Ding and Dokholyan, 2006). To construct ensemble of simulated “orthologs”, we first primed a population by evolving it until it reached a state of dynamic mutation-selection balance (Goyal *et al*., 2012). We then used the output to perform 160 independent evolutionary simulations under selection for folding stability. We ran each simulation for 700,000 generations, ensuring that the distribution of pairwise sequence identities for simulated proteins matches that of the orthologs. For both sets, the distribution is a Gaussian centered around 36% pairwise identity. Because each run produces as many sequences as there are cells in the population, we narrowed down our set by randomly sampling 5 sequences from each run for a total of 800 simulated DHFR sequences. For real orthologous sequences, we retrieved the top 250 hits of a protein BLAST for the 192 amino acid *Candia albicans* dihydrofolate reductase (DHFR), from which we excluded sequences longer than 220 bp. We used the 163 remaining sequences to construct a multiple sequence alignment using Clustal Omega (Sievers *et al*., 2011). To compare sequence conservation, we used the Kullback-Leibler conservation score, which is a measure of relative entropy (Kullback and Leibler 1951) for each residue *z*:

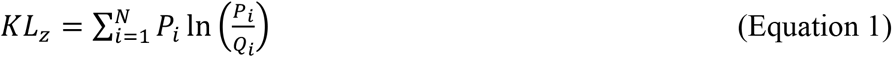

where *P_i_* is the observed frequency of amino acid *i* in that specific residue and *Q_i_* is the background natural frequency of that specific amino acid shared amongst residues in orthologs. A higher KL score implies a higher conservation of that residue’s identity throughout evolution. Conversely, when KL is closer to zero, that residue’s identity is frequently substituted. Because thermodynamic stability is the major evolutionary pressure on DHFR, our computational model should be able to recapitulate the pattern of native sequence conservation. Indeed, as shown in Figure 4, the sequence conservation of simulated DHFR sequences is significantly correlated with real DHFR orthologs.

**Figure 4.**
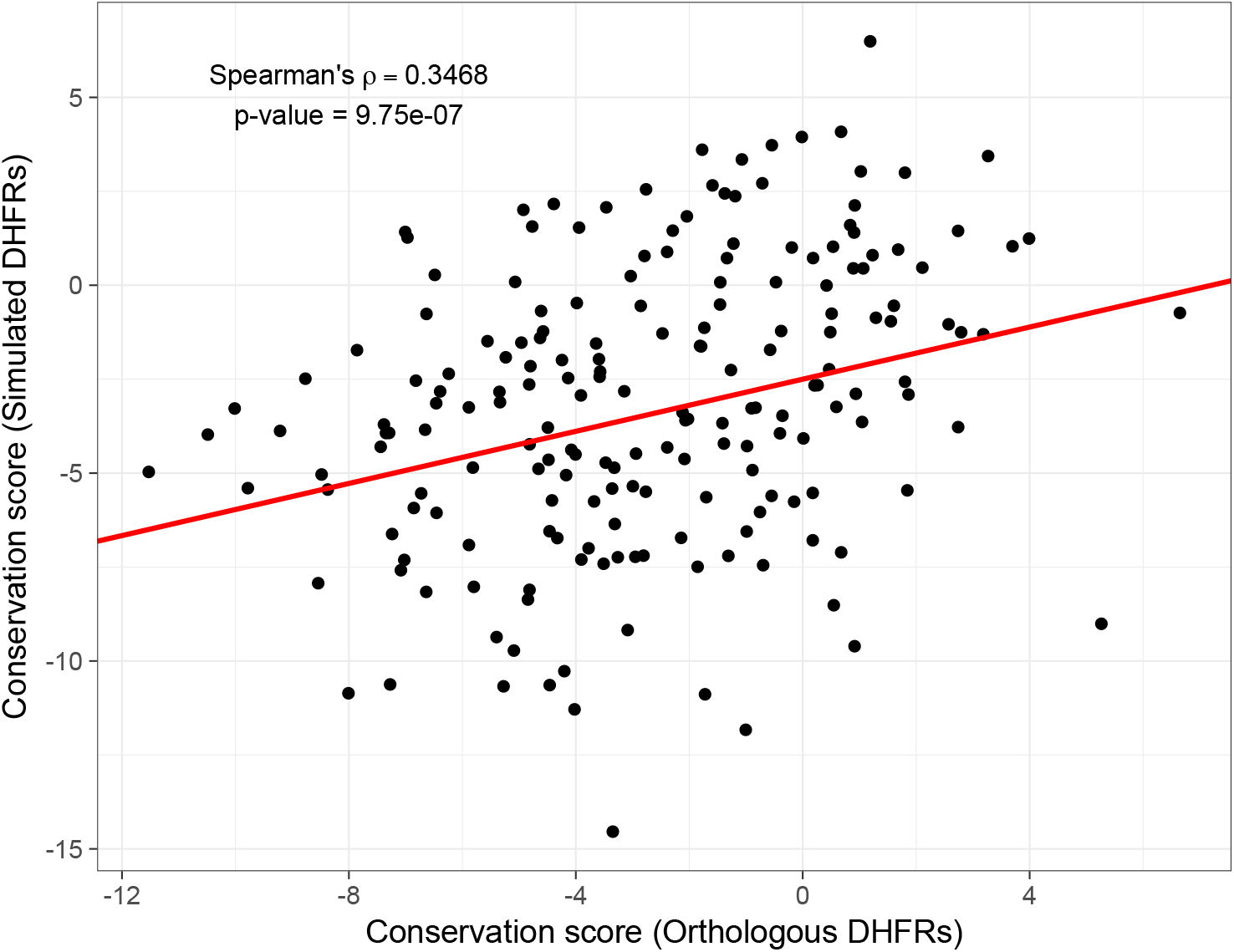
Test case 2: evolution under selection for thermodynamic stability. SodaPop captures a significant fraction of sequence conservation in DHFR.

### Performance and runtime

SodaPop is the first publicly available tool which can effectively simulate multi-scale molecular evolution and polyclonal population dynamics in an all-encompassing framework. We benchmarked SodaPop for multiple population sizes and number of generations. All simulations were run on a standard iMac desktop with a 3.2GHz Intel Core i5 processor and 16GB memory. Figure 5 shows that runtime is quasi-monomial with respect to population size. We limited our desktop benchmarking to N=10^6^ cells, as higher orders of magnitude induce a shift in performance due to a RAM bottleneck. Explicit simulation of populations with higher orders of magnitude requires a larger amount of memory than the current standard in commercial desktop computers. Larger population sizes can be simulated on high-performance computing clusters where memory allocation is not limiting. However, simulating up to a million cells for long time periods is entirely tractable using standard desktop computers.

**Figure 5.**
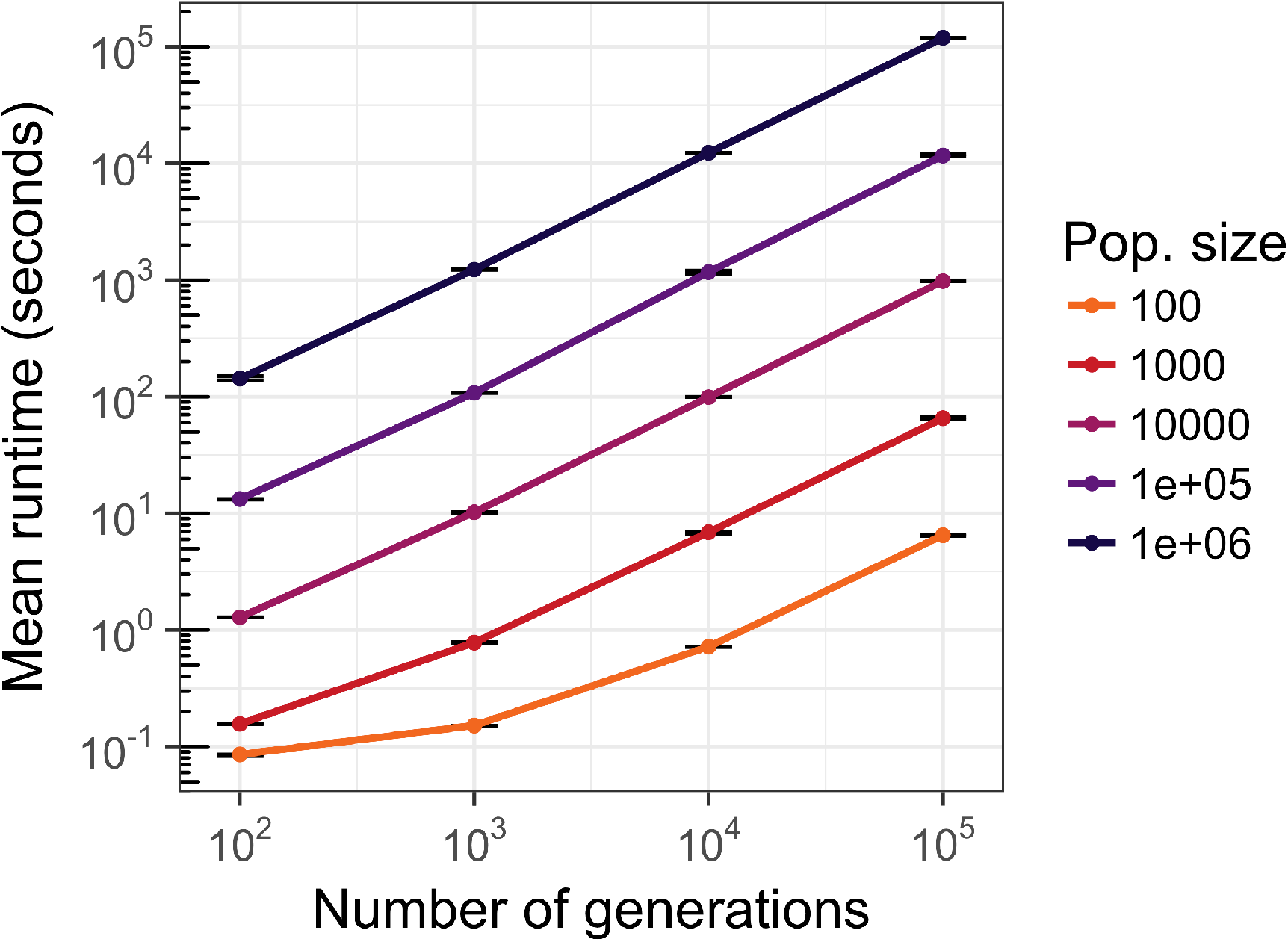
Benchmarking of SodaPop for different population sizes. Runtime of SodaPop with varying population sizes and simulation length. The time step for each test case was set to 0.01 N.

## Conclusion

There are several features that can reduce the required memory for the performance of SodaPop. First, using a binary encoding of the genetic code should reduce the memory required to store a single cell by a significant factor without incurring any information loss. Second, collapsing lineages within a single consensus sequence could also reduce the memory load, at a cost of information loss. These are currently under development for future versions. Considering the need to address questions at the interface of molecular evolution and population genetics, and with most of the current computational methods unable to account for explicit clonal dynamics, we believe SodaPop provides a comprehensive and extensible framework that can encompass a wide array of evolutionary scenarios.

## Acknowledgements

We would like to thank members of the Serohijos lab, especially Pouria Dasmeh, Ahn-Tien Ton and Christopher Savoie for testing the program and contributing valuable questions and ideas.

